# Soluble cyanobacterial carotenoprotein as a robust antioxidant nanocarrier and delivery module

**DOI:** 10.1101/823880

**Authors:** Eugene G. Maksimov, Alexey V. Zamaraev, Evgenia Yu. Parshina, Yury B. Slonimskiy, Tatiana A. Slastnikova, Alibek A. Abdrakhmanov, Pavel A. Babaev, Svetlana S. Efimova, Olga S. Ostroumova, Alexey V. Stepanov, Anastasia V. Ryabova, Thomas Friedrich, Nikolai N. Sluchanko

## Abstract

To counteract oxidative stress, antioxidants including carotenoids are highly promising, yet their exploitation is drastically limited by the poor bioavailability and fast photodestruction, whereas current delivery systems are far from being efficient. Here we demonstrate that the recently discovered nanometer-sized water-soluble carotenoprotein from *Anabaena* (termed CTDH) transiently interacts with liposomes to efficiently extract carotenoids via carotenoid-mediated homodimerization, yielding violet-purple protein samples amenable to lyophilization and long-term storage. We characterize spectroscopic properties of the pigment-protein complexes and thermodynamics of liposome-protein carotenoid transfer and demonstrate the highly efficient delivery of echinenone form CTDH into liposomes. Most importantly, we show carotenoid delivery to membranes of mammalian cells, which provides protection from reactive oxygen species. The described carotenoprotein may be considered as part of modular systems for the targeted antioxidant delivery.

**Significance statement:** Carotenoids are excellent natural antioxidants but their delivery to vulnerable cells is challenging due to their hydrophobic nature and susceptibility to degradation. Thus, systems securing antioxidant stability and facilitating targeted delivery are of great interest for the design of medical agents. In this work, we have demonstrated that soluble cyanobacterial carotenoprotein can deliver echinenone into membranes of liposomes and mammalian cells with almost 70 % efficiency, which alleviates the induced oxidative stress. Our findings warrant the robustness of the protein-based carotenoid delivery for studies of carotenoid activities and effects on cell models.

## Introduction

Formation of reactive oxygen species (ROS) accompanies electron transfer during aerobic respiration or photosynthesis. Since high ROS levels may be harmful to cells, antioxidants are crucial for maintaining their normal functioning (1, 2). Carotenoids are natural antioxidants playing important roles in photoprotection and regulation of photosynthetic activity of higher plants, algae, and cyanobacteria. Due to the very short lifetime of the excited state (3), carotenoids acting as excitation energy acceptors can rapidly convert light energy into heat, thereby reducing the probability of ROS formation. Mammalian cells cannot produce carotenoids, but some types are vitally needed not only as antioxidants. For example, β-carotene is a source of retinal, the cofactor of visual photoreceptors (4). Alongside with the reported anti-cancer (5), anti-tumor or anti-dermatosis abilities of carotenoids (6, 7), numerous studies revealed effects of carotenoids in human chronic diseases including the so-called canthaxanthin retinopathy, retinal dystrophy or aplastic anemia (8). In any case, from visual pigments to coloration of bird feathers, carotenoids come from diet and must be delivered to specific tissues and cells to perform their functions (9). While being transported by blood, nutritional carotenoids most often are found in lipoproteins harboring the sites suitable for promiscuous binding of different lipophilic molecules (10). However, the mechanisms which allow for the delivery of carotenoids into cells in a specific and targeted manner are unknown.

Modern strategies to deliver carotenoids into tissues are based on liposomes, niosomes, solid lipid nanoparticles, polysaccharides and oligosaccharides inclusion complexes which can be constructed in a controlled manner (11–15). The fusion of liposomes pre-loaded by carotenoids with cellular membranes causes delivery of the antioxidants into different cell compartments. However, the efficiency of carotenoid uptake by cells is limited, reportedly, incubation of cells with liposomes with micromolar carotenoid concentrations results in picomolar concentrations in cells (11). This is further complicated by poor carotenoid stability and fast photodestruction. For targeted delivery, a conjugation with antibodies specific to some cell surface components may be necessary. Alternative protein-based modular constructions are being intensively developed since protein sequence and functionality can be effectively engineered (16–18). Fortunately, natural water-soluble carotenoid-binding proteins may provide the best opportunities for carotenoid transportation and targeted delivery.

The structures of the photoactive Orange Carotenoid Protein (OCP) and some of its recently discovered homologs are optimized by evolution to ensure carotenoid retrieval from the membrane (19–21), since their functionality requires delivery of the carotenoid molecule to the antenna complexes to quench overexcitation (22). Besides that, OCP is also an efficient ROS quencher (23). Upon expression in carotenoid-producing *Escherichia coli* strains, OCP-like proteins can bind echinenone (ECN), canthaxanthin (CAN), zeaxanthin and β-carotene (24–26). Assembly of these water-soluble carotenoproteins requires carotenoid absorption from membranes, however, the mechanism of this process is barely known. Very recently, it was demonstrated *in vitro* that a natural homolog of the C-terminal domain of OCP (hereinafter CTDH) from *Anabaena* can take keto-carotenoids from membranes of overproducing *E.coli* strains (27, 28). This leads to the maturation of the initially colorless CTDH apoprotein into a violet, soluble nm-sized holoprotein via the carotenoid-mediated protein homodimerization (24, 28).

To avoid the heterogeneity associated with the uncontrolled lipid, carotenoid, and protein content of the *E.coli* membranes used in the previous work (27), we employed simpler model and used liposomes to study the assembly of a water-soluble carotenoid carrier. In contrast to our initial expectations, we found that carotenoid transfer between the membrane and the protein is reversible and that the efficiency of this process critically depends on particular protein-membrane and protein-carotenoid interactions. Furthermore, we demonstrate that soluble cyanobacterial carotenoid carriers can be used for the delivery of carotenoids into mammalian cells and provide protection from ROS. We discuss the outreach of such approaches for biomedical applications.

### Direct interactions of ApoCTDH with the membrane

In order to increase the probability of carotenoid uptake, water-soluble ApoCTDH should directly interact with the lipid bilayer, which was tested in a series of experiments with different model membranes.

The threshold voltages (*V_bd_*) that caused the electrical breakdown of the DPhPC and DOPC membranes in the absence of ApoCTDH were 460 ± 40 mV and 370 ± 20 mV, respectively (*Figure 1A*). The introduction of ApoCTDH into membrane bathing solution up to a concentration of 5 μM led to about twofold decrease in *V_bd_* of DPhPC-bilayers (230 ± 15 mV). For DOPC-bilayers, *V_bd_* decreased by 1.5 times (down to 225 ± 15 mV), suggesting that ApoCTDH interaction decreases the electrical stability of model membranes. In contrast, the addition of ApoCTDH did not affect the steady-state conductance of DPhPC- and DOPC-bilayers induced by K^+^-nonactin. This indicates that the distribution of the electrical potential on the membrane/water interface (membrane boundary potential, φ_*b*_) remains unchanged upon protein adsorption (Δ φ_*b*_ = 1 ± 1 mV). Thus, interactions of ApoCTDH and membranes are transient but detectable, which is reasonable, considering the possible functional role of this protein as a soluble carotenoid carrier.

**Fig. 1.**
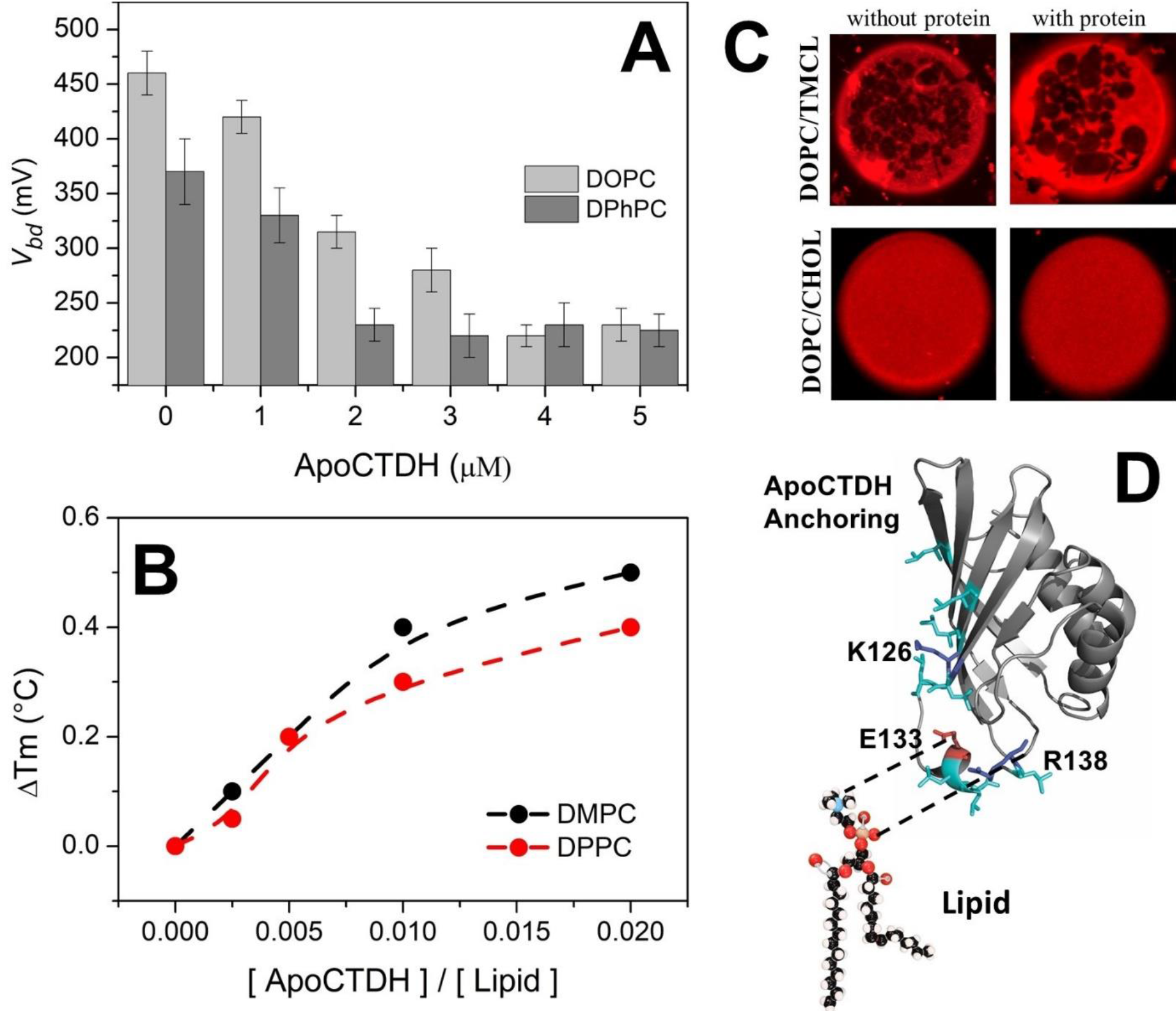
Interactions of ApoCTDH with membranes. (**A**) – threshold voltages (*V_bd_*) that caused electrical breakdown of DOPC and DPhPC membranes at different concentration of ApoCTDH in solution. (**B**) – the increase of phase transition temperature of liposomes composed of DMPC and DPPC in relation to the ApoCTDH concentration. (**C**) - fluorescence micrographs of giant unilamellar vesicles membranes made from 50 mol % DOPC and 50 mol % TMCL (top row) and 67 mol % DOPC and 33 mol % CHOL (bottom row) in the absence and in the presence of ApoCTDH. The lipid:protein ratio was 100:1. The liquid, disordered phase appears red, while the solid, ordered phase remains dark. Image size is 15×15 μm. (**D**) – model of ApoCTDH interactions with PC. Glu-133 is shown in red, positively charged residues are shown in blue, leucines are shown in cyan. The structure of ApoCTDH is drawn in Pymol using PDB ID 6FEJ chain A (28).

Using confocal microscopy, the lateral heterogeneity of vesicle membranes formed from DOPC/DPPC, DOPC/DMPC, DOPC/SM and DOPC/TMCL (each 50/50 mol %) before and after addition of ApoCTDH into liposome suspension was studied. In these lipid systems, the coexistence of the liquid-disordered and gel-lipid phases (*l_d_* + *s_o_*) was observed. Micrographs (*Figure 1C*) demonstrate that addition of ApoCTDH at a 100:1 molar ratio did not promote the induction of visible ordered domains of micro sizes and did not change the shape of lipid vesicles and the phase segregation scenario.

Although visible phase segregation was not affected by interactions of ApoCTDH with membranes, an appreciable increase of the phase transition temperature (*Figure 1B*) was found. For liposomes composed of DMPC, the phase transition occurs at 22.8 °C. Addition of protein up to a lipid:protein ratio of 100:1 causes an increase of the phase transition temperature by ~0.5 °C. A width at half-height of the main peak in the endotherm (*T_1/2_*), informing about cooperativity of transition, was not changed. The similar results were obtained with DPPC (*Figure 1B*), which proves that that interaction of the protein with the membrane does not depend on the thickness of its hydrocarbon core. This suggests that ApoCTDH likely interacts with the polar heads of neighboring membrane lipids in the model membranes owing to electrostatic interactions. Considering the ApoCTDH structure (28), it is probable that the polymorphous C-terminal tail (CTT), featuring Glu-133 and Arg-138 residues, can act as an anchor to increase the probability of carotenoid uptake (*Figure 1D*). Several Leu residues may also contribute to the latter process.

Our results are in excellent agreement with the fact that the CTT absence reduces the rate of holoprotein formation (28). Considering also that in the crystallographic ApoCTDH dimer, the CTT partially blocks the so-called carotenoid tunnel and adopts different conformations, it is likely that this structurally mobile Leu-rich motif plays a critical role in various carotenoid transfer processes.

### Carotenoid uptake by Apo-CTDH from liposomes

Further, we assessed the optical response of ECN and CAN in different environments (SI, *Figure S1*), which provided us with the spectroscopic signatures to study interactions of liposomes loaded with carotenoids and ApoCTDH (*Figure 2*).

**Fig. 2.**
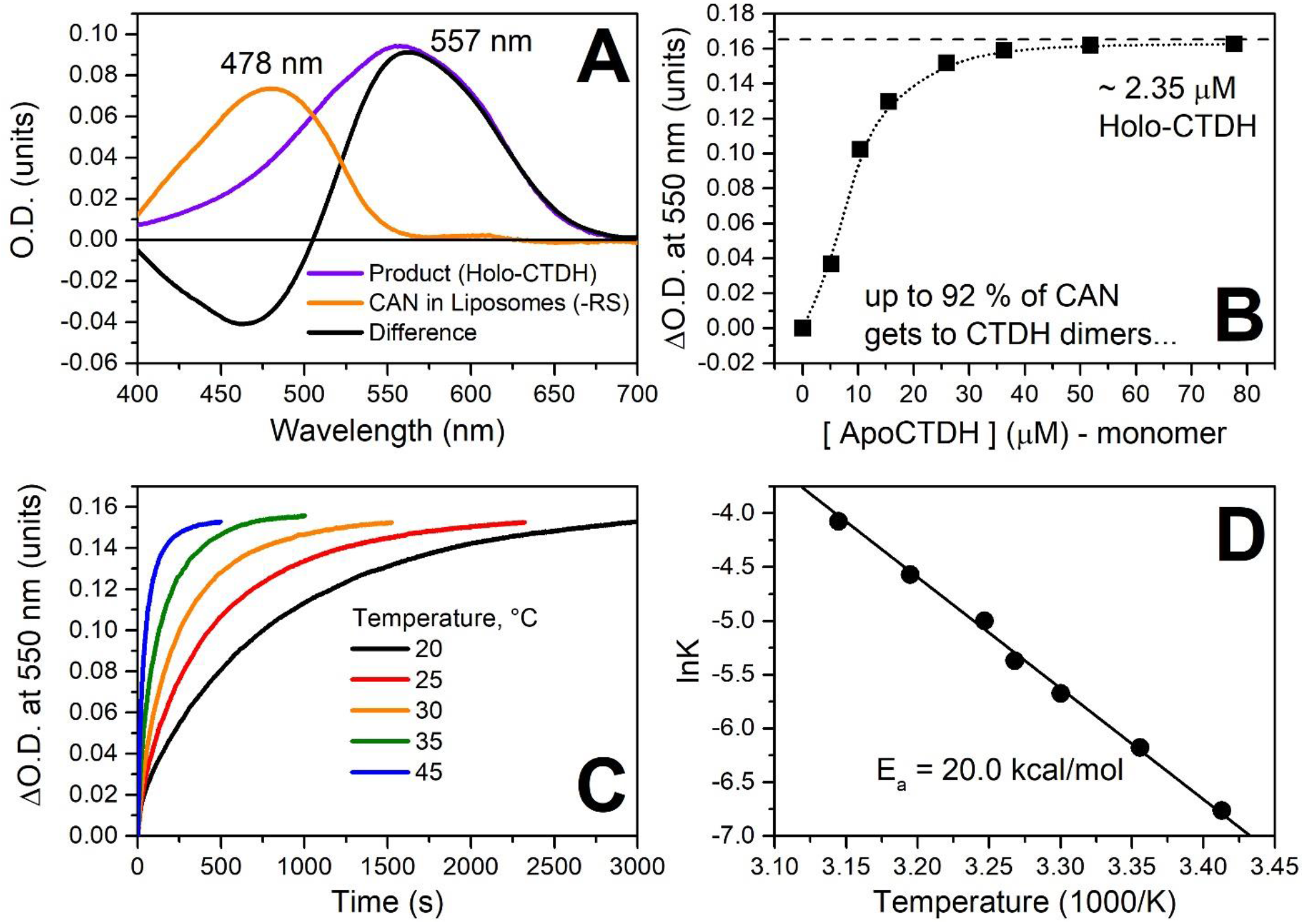
CAN uptake by ApoCTDH from liposomes. (**A**) – absorption spectra of CAN in liposomes and after incorporation into CTDH. Rayleigh scattering (RS) was subtracted. (**B**) – increase of O.D. at 550 nm upon addition of different amounts of ApoCTDH to CAN-containing liposomes. (**C**) – time-courses of accumulation of HoloCTDH due to carotenoid uptake from liposomes at different temperatures and (**D**) – the corresponding temperature dependency of rate constants (Arrhenius plot).

After addition of colorless ApoCTDH to yellowish liposomes containing CAN, the solution turned violet, with the concomitant pronounced spectral changes (*Figure 2A*). This indicated that CTDH interacts with membranes to efficiently extract membrane-embedded carotenoids and homodimerize. The rate of this process depended on temperature (*E_A_* ~20.0 kcal/mol) (*Figure 2C and D*). Titration experiment monitoring the increase in 550 nm absorption yielded saturation at ten ApoCTDH monomers per single carotenoid molecule (*Figure 2B*). At such ratios, almost all carotenoids were extracted from liposomes, giving up to 92% of violet CTDH(CAN). In contrast, only ~ 35% of carotenoid was extracted by CTDH from ECN-containing liposomes (*Figure S2*). Accumulation of CTDH(ECN) was almost 6 times slower comparing to CTDH(CAN) at 30 °C and similar protein concentration.

### Carotenoid delivery into liposomes

Since ~65% ECN remains in liposomes even in the ApoCTDH excess, one can expect the equilibrium between carotenoid uptake from and carotenoid delivery into membranes. In theory, the equilibrium can be shifted, and the delivery might be more efficient, which prompted us to test different carotenoprotein holoforms as carotenoid donors for the liposomes. Upon mixing of CAN-containing COCP (29), OCP^AA^ (30), RCP (20), and CTDH with liposomes, we failed to observe any substantial carotenoid transfer. In contrast, the addition of CTDH with embedded ECN led to a decrease of 530 nm absorption and a concomitant increase of 440 nm absorption (*Figure 3A*). This can be followed visually, by a color change from violet-purple to light yellow, and directly confirmed by size-exclusion spectrochromatography (*Figure S3*), together indicating productive carotenoid delivery from the protein to liposomes. Upon titration by liposomes, we reached saturation corresponding to a ~ 70 % efficiency of carotenoid delivery (*Figure 3B*).

**Fig. 3.**
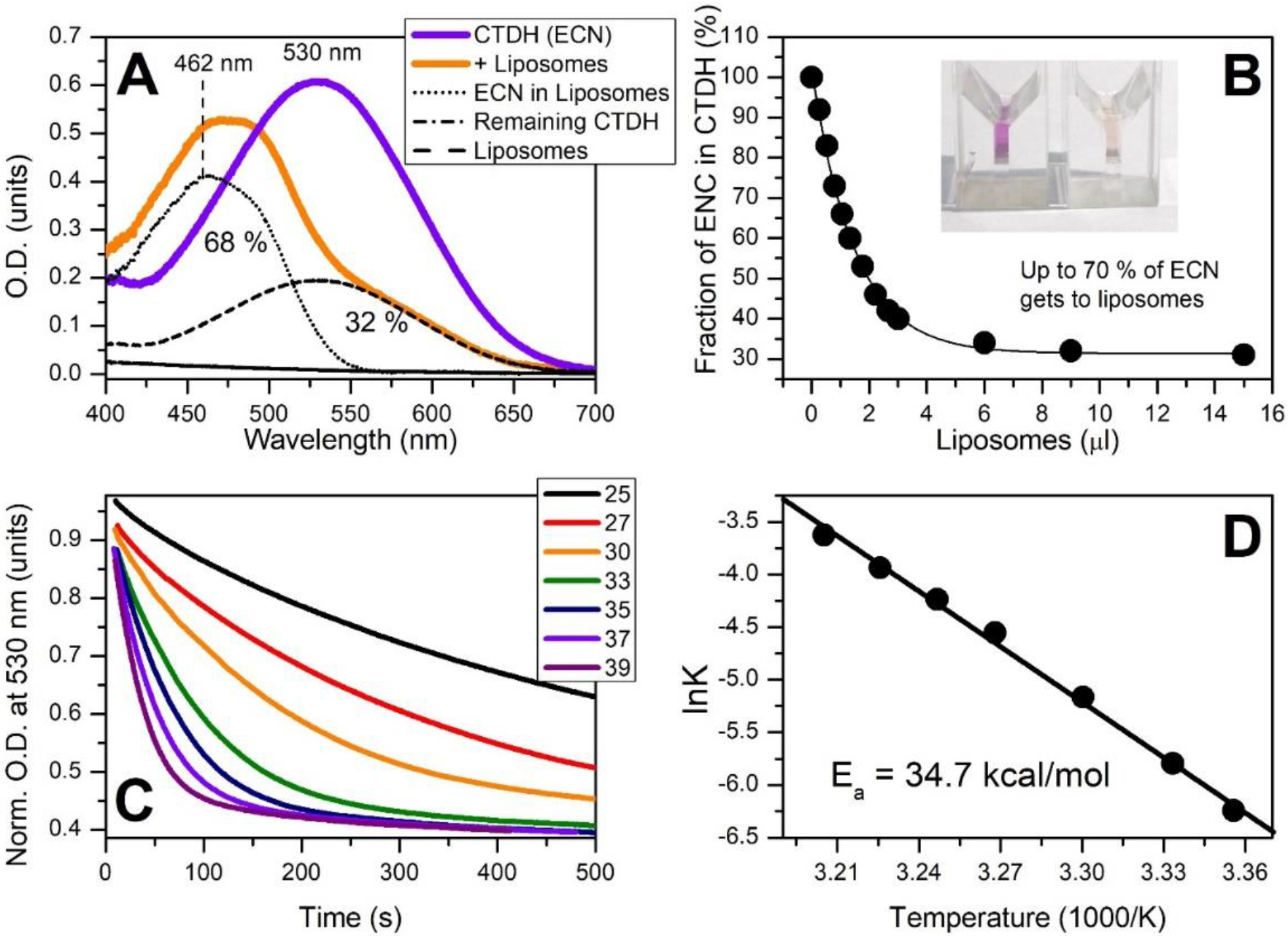
ECN delivery by CTDH into liposomes. (**A**) – the absorption spectrum of CTDH (ECN) before (purple) and after addition of liposomes (orange). Percentage of carotenoid remaining in CTDH after addition of liposomes was determined by the resulting spectrum decomposition into CTDH(ECN) absorption and scattering of liposomes, considering also the fact that ECN absorption in liposomes is significantly blue-shifted and does not overlap with protein-bound ECN in the 550–700 nm region. (**B**) – percentage of carotenoid remaining in CTDH upon the increase of concentration of liposomes in solution. The dependency was approximated by exponential function in order to determine the maximum yield of carotenoid delivery. The inset shows cuvettes with CTDH(ECN) solution before (left) and after incubation with liposomes. (**C**) - time-courses of optical density at 530 nm corresponding to disappearance of holo-CTDH due to carotenoid delivery into liposomes at different temperatures and (**D**) – the corresponding temperature dependence of rate constants (Arrhenius plot).

The rate of ECN delivery into liposomes by CTDH was very sensitive to temperature, yielding *E*_A_ of ~34.7 kcal/mol (*Figure 3C and D*), which is larger than the energy barrier for CAN uptake by ApoCTDH and suggests significant rearrangements of the CTDH conformation upon carotenoid release. Noteworthy, the ECN delivery rate is significantly higher than the rate of ECN uptake at 30 °C (*Figure S2*) and could be even faster at 37°C, providing a solid thermodynamic foundation for virtually unidirectional ECN delivery at physiological temperatures.

### Carotenoid delivery by CTDH into mammalian cells

We next questioned whether CTDH-mediated ECN delivery can occur with more complex, biomedically relevant membrane models.

After incubation of HEK293, HeLa, neuroblastoma (Tet21N), and ovary carcinoma cell suspensions in the presence of CTDH(ECN), characteristic changes of carotenoid color from purple-violet into yellow (*Figure 4*) were observed, pointing to ECN migration into cell membranes. Measurements of ECN absorption in eukaryotic cell lines are complicated due to significant light scattering. To circumvent this difficulty, characteristic Raman signatures (see *Figure S1* and description in SI) were used to study carotenoid delivery and distribution in cells. *Figure 4A* shows that the Raman spectrum of CTDH(ECN) changes after incubation with liposomes: the *ν*_1_ band becomes significantly broader due to the contributions from two different fractions of ECN: one embedded in CTDH (~ 30 %), while the rest residing in membranes. The same distribution was observed upon incubation of HeLa cells in the presence of CTDH(ECN) (data not shown). After washing out the residual CTDH with fresh culture medium we were able to analyze intracellular carotenoid distribution by Raman microscopy. After incubation with CTDH(ECN), normally carotenoid-free HeLa cells demonstrated characteristic spectral signatures of ECN in membranes, while the contribution from CTDH(ECN) completely vanished. Using the microscope, we found that Raman signatures of ECN colocalized with the cells (*Figure 4B*). Notably, the distribution of carotenoids across the cell, which can be estimated by the intensity of *ν*_1_ band, is not homogeneous (*Figure 4C*). Interestingly, CTDH fusion with the red fluorescent protein (RFP) allowing imaging, can bind ECN from different sources (proteins or liposomes), but does not localize in cells (*Figure S5*) excluding protein adsorption or endocytosis as the reason for increased carotenoid content of cells. Based on these observations, we conclude that CTDH(ECN) approaches the outer cell membrane, unloads the carotenoid, and remains outside of the cell, while carotenoid is redistributed across cellular membranes due to other, internal transport mechanisms.

**Fig. 4.**
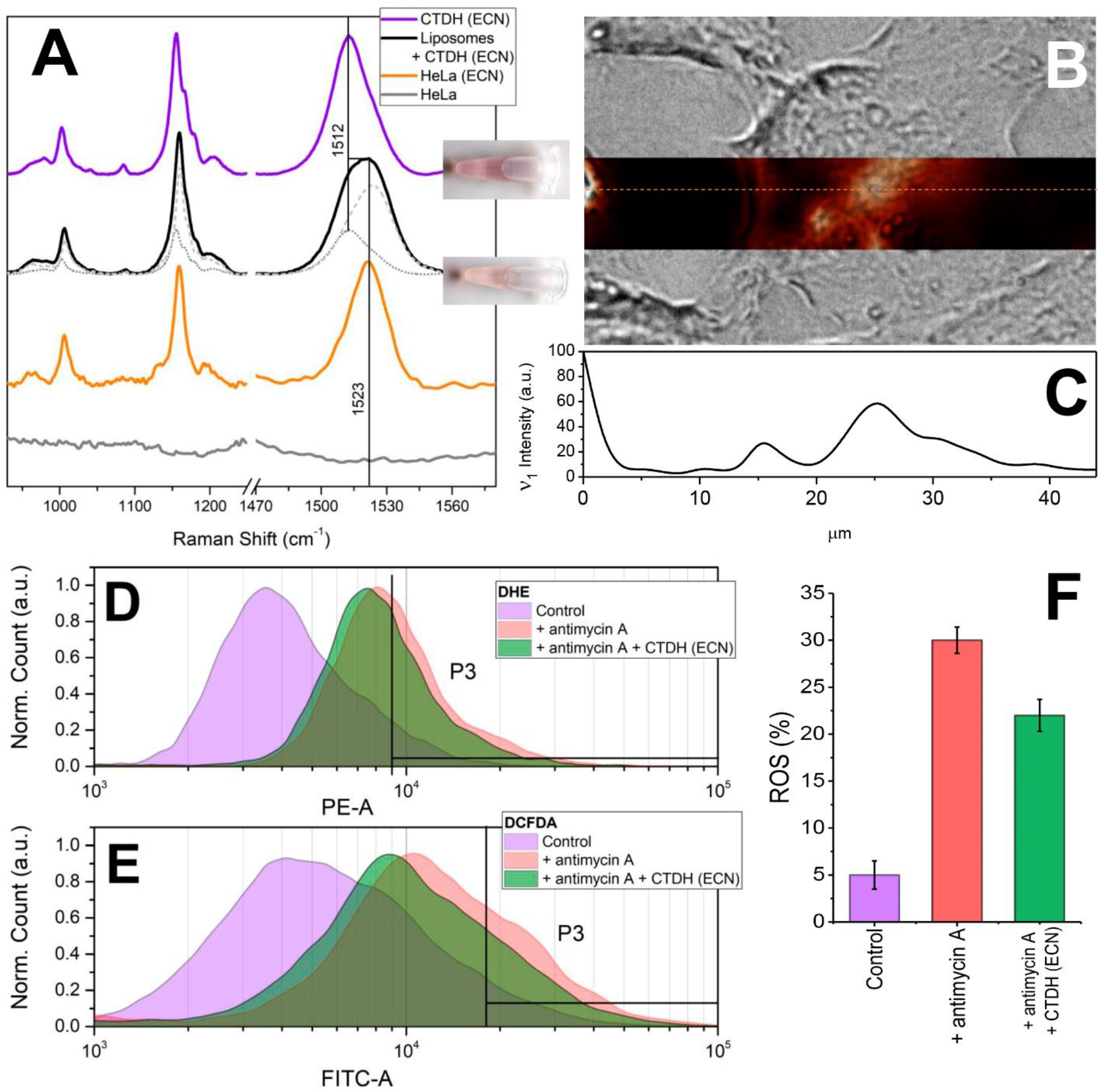
Carotenoid delivery to mammalian cells. (**A**) - Raman spectra of ECN in CTDH (violet) and in liposome suspensions after incubation with CTDH(ECN) (black). The spectrum was decomposed into a linear combination of Raman spectra of ECN in CTDH (30 %) and in liposomes (70 %) (see *Figure S1*). The orange line shows characteristic Raman spectra of ECN in HeLa cells. The grey line shows Raman of HeLa cells before incubation with CTDH (ECN), no bands in the carotenoid region were observed. Inset shows a tube with a suspension of HeLa cells containing 5 μM of CTDH(ECN) before (top) and after two hours of incubation (bottom) at 37 °C. (**B**) – overlay of microscopic images of a typical HeLa cell in transmitted light with the of *ν*_1_ band intensity (at 1522 cm^-1^ minus background at 1550 cm^-1^) presented in pseudo colors. Dotted line shows a cross-section through the Raman image, the corresponding distribution of *ν*_1_ band intensity is shown in (**C**). Effect of carotenoid delivery by CTDH (ECN) into Tet21N cells on ROS production induced by antimycin A was determined by DHE (**D**) and DCFDA (**E**) staining and flow cytometry. (**F**) – the yield of ROS in Tet21N cells (purple, control sample), under oxidative stress induced by antimycin A treatment (red) and in cells pre-incubated with CTDH (ECN) and treated by antimycin A (green).

### Carotenoid delivery to mammalian cells alleviates oxidative stress

Having found that ECN could be delivered to plasmalemma, we tested if this natural antioxidant can reach mitochondria and counteract oxidative stress. Using dihydroethidium (DHE) and 2′-7′-dichlorodihydrofluorescein diacetate (DCFDA) staining protocols we analyzed ROS accumulation by flow cytometry. The addition of antimycin A (31) to Tet21N cell line induces ROS production from 5 up to 30%. As a positive antioxidant control, we used *N*-acetylcysteine (NAC) which has a free radical scavenging property and almost completely prevents the accumulation of ROS after the antimycin A treatment in Tet21N cells (32). Incubation of cells in the presence of 1 μM CTDH(ECN) decreased ROS production by 25% (from 30% to 22%) in both types of experiments (DHE and DCFDA staining) (*Figure 4*). Thus, ECN delivered into mammalian cells by the cyanobacterial protein can protect it from the oxidative stress.

### Conclusions and perspectives

We found that ApoCTDH transiently interacts with the membrane, which appears a critical step for the formation of water-soluble carotenoid holo-proteins. Comparing the optical response of CTDH with different embedded carotenoids (ECN or CAN) we have found that the stability of protein-carotenoid complex strongly depends on the presence of hydrogen bonds between the keto-group of the carotenoid and conserved aromatic residues, subject to engineering in order to modulate the carotenoid binding efficiency. The relative stability of such protein-chromophore interactions determines the ability of the system to take, transport and deliver carotenoids from lipid membranes into other compartments. We assume that the protein-chromophore interactions that allow CTDH to bind carotenoids lacking keto-groups in one (ECN) or both β-rings (like β-carotene) are relatively weak as holo-forms appear only at large protein excess. We scrutinized the process of carotenoid uptake from membranes by CTDH and showed that ECN distribution is in a dynamic equilibrium which is shifted from the protein to membrane (35 vs. 65 %, respectively), permitting efficient delivery of carotenoids into membranes (*Figure 5*). Moreover, light could be potentially used in order to activate the process of carotenoid delivery into membranes by CTDH from the photoconvertible OCP (*Figure S4*) (24).

**Fig. 5.**
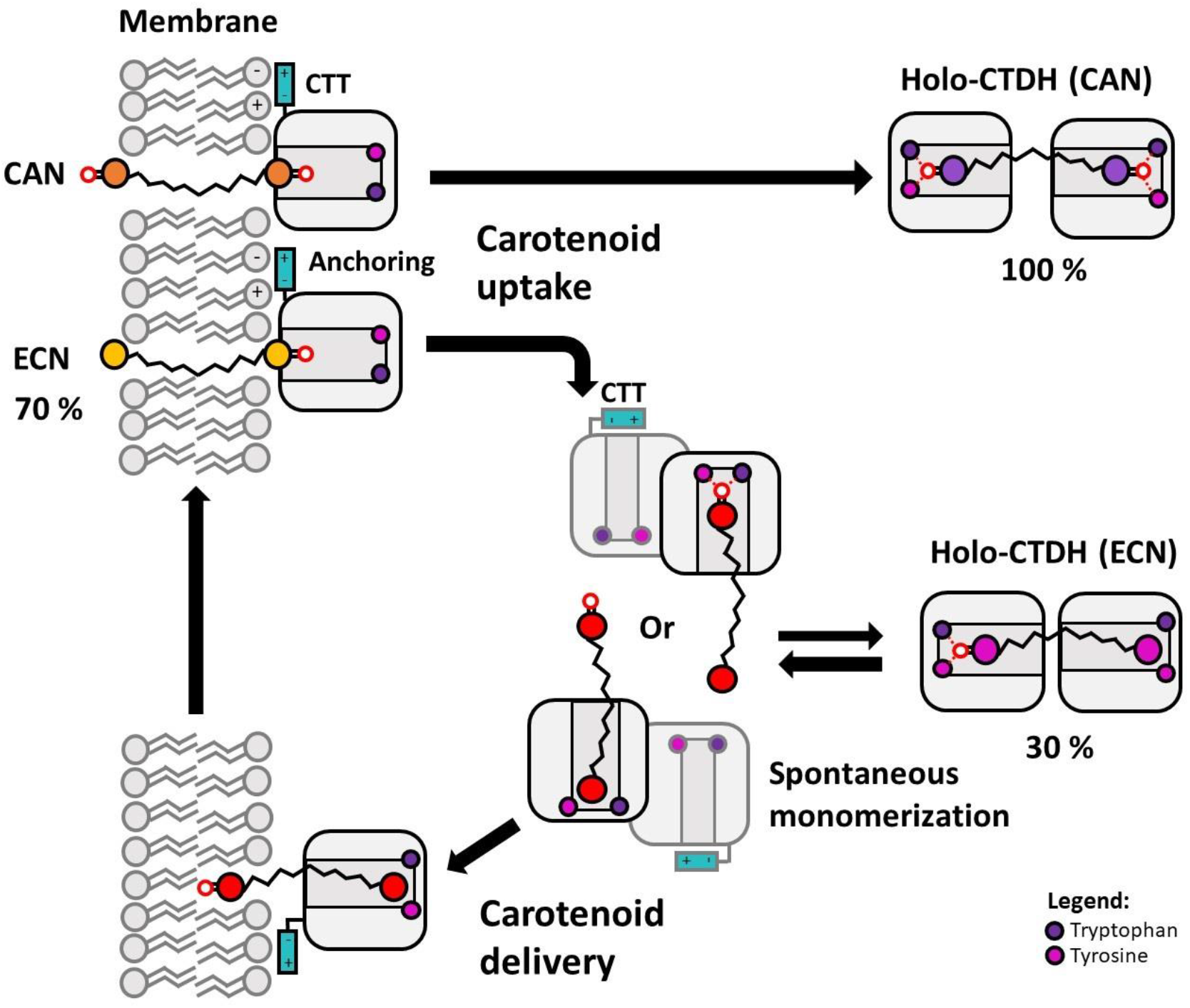
Proposed model for CTDH-mediated carotenoid uptake and delivery. Carotenoid uptake by CTDH from the membrane is promoted by electrostatic interactions of the CTT and lipid head groups resulting in anchoring and formation of a transient complex between the membrane and protein facing carotenoid binding cavity towards the membrane. In such complex, spontaneous translocation of carotenoid into hydrophobic part of the protein may be stabilized by formation of the hydrogen bond between carotenoid keto-group and Trp/Tyr. Due to a significant length of the carotenoid molecule it requires two CTDH subunits to isolate it from solvent. The presence of two keto-groups in CAN results in the most efficient carotenoid binding in the CTDH dimer, while ECN binding is apparently weaker. Since both types of CTDH can transfer carotenoids into other proteins we postulate that spontaneous monomerization of protein dimer occurs regardless of the carotenoid type (24). However, only CTDH monomers in which keto-group of ECN loses connection with the protein give carotenoid an opportunity to escape another protein subunit and get to the membrane once again.

Carotenoids are excellent natural antioxidants but their delivery to vulnerable cells is challenging due to their hydrophobic nature and susceptibility to degradation. Thus, systems securing antioxidant stability and facilitating targeted delivery are of great interest for the design of medical agents (13, 15, 33–37). In this work, we have demonstrated that CTDH can deliver ECN into membranes of liposomes and mammalian cells with almost 70 % efficiency, which, in Tet21N cells, alleviates the induced oxidative stress. Our findings warrant the robustness of the protein-based carotenoid delivery for studies of carotenoid activities and effects on cell models. Alongside with the unseen delivery efficiency, the remarkable stability of OCP-like proteins and rapid carotenoid release rates (minutes) comparing to liposomes (days (11)), the greatest advantage is the ability to construct genetically encoded modular systems exploiting a toolbox of different functional modules. In this regard, use of cyanobacterial water-soluble proteins seems encouraging for numerous biomedical applications and can benefit from their tolerance to lyophilization and astonishingly long shelf-life (years). Last but not least, the ability of CTDH to extract CAN from membranes could potentially be utilized for reversing conditions like canthaxanthin retinopathy associated with the ill accumulation of this dietary carotenoid in tissues.

## Materials and Methods

### Materials

Synthetic 1,2-diphytanoyl-sn-glycero-3-phosphocholine (DPhPC), 1,2-dioleoyl-sn-glycero-3-phosphocholine (DOPC), 1,2-dipalmitoyl-sn-glycero-3-phosphocholine (DPPC), 1,2-dimyristoyl-sn-glycero-3-phosphocholine (DMPC), sphingomyelin (Brain, Porcine) (SM), 1’,3’-bis[1,2-dimyristoyl-sn-glycero-3-phospho]-sn-glycerol (sodium salt) (TMCL), cholesterol (CHOL) and 1,2-dipalmitoyl-sn-glycero-3-phosphoethanolamine-N-(lissamine rhodamine B sulfonyl) (Rh-DPPE) were obtained from Avanti Polar Lipids, Inc. (Pelham, AL). Nonactin, NaCl, NaOH, HEPES, EDTA, and hexadecane were purchased from Sigma Chemical (St. Louis, MO, USA).

### Cloning, protein expression, and purification

The identity of the constructs and the presence of mutations were verified by DNA sequencing (Evrogen, Moscow, Russia). The obtained plasmids were used to transform chemically competent cells. Proteins were expressed using induction by 1 mM isopropyl-β-thiogalactoside in the presence of kanamycin and ampicillin.

Holoforms of CTDH were expressed in echinenone (ECN) and canthaxanthin (CAN)-producing *Escherichia coli* cells essentially as described earlier (38). All His_6_-tagged proteins were purified by immobilized metal-affinity and size-exclusion chromatography to electrophoretic homogeneity and stored at 4 °C in the presence of 3 mM sodium azide. Protein concentrations were determined at 280 nm using calculated protein-specific molar extinction coefficients.

### Lipid bilayer setup, recording system, and calculations

Virtually solvent-free planar lipid bilayers were prepared using a monolayer-opposition technique (39) on a 50-*μ*m-diameter aperture in a 10-*μ*m-thick Teflon film separating two (*cis*- and *trans*-) compartments of a Teflon chamber. The aperture was pretreated with hexadecane. Lipid bilayers were made from pure DOPC or pure DPhPC. Solutions of 0.1 M NaCl, 1 mM EDTA were buffered by 5 mM HEPES-NaOH at pH 7.4. After the membrane was completely formed and stabilized, protein AnaCTDH-apo was added to the *cis*-compartment from a stock solution in the buffer to obtain final concentration ranging from 1 to 5 μM. Ag/AgCl electrodes with 1.5% agarose/2 M KCl bridges were used to apply the transmembrane voltage (*V*) and measure the transmembrane current. “Positive voltage” refers to the case in which the *cis*-side compartment is positive with respect to the *trans*-side.

Current measurements were carried out using Axopatch 200B amplifier (Axon Instruments) in the voltage clamp mode. Data were digitized by Digidata 1440A and analyzed using pClamp 10 (Axon Instruments) and Origin 8.0 (Origin Lab). Data acquisition was performed with a 5 kHz sampling frequency and low-pass filtering at 100 Hz. The current tracks were processed through an 8-pole Bessel 100-kHz filter.

The threshold voltages that cause pure DOPC (or pure DPhPC) membrane electrical breakdown before and after addition of AnaCTDH-apo protein into bathing solution up to 1-5 μM, *V_bd_*, were measured using triangle-shaped ramps (±10 mV/s) in the range of 0 to ±*V_bd_*. No difference between positive and negative voltages was observed.

### Measurement of the membrane boundary potential

The steady-state conductance of K^+^-nonactin was modulated via the one-sided addition of protein ApoCTDH from 140 mM stock solutions in buffer to the membrane-bathing solution (0.1 M NaCl, 1 mM EDTA, 5 mM HEPES-NaOH, pH 7.4) to obtain a final threshold concentrations defined by the electrophysiological methods that the compounds increase the ion permeability of the lipid bilayer. The membranes were composed of DOPC or DPhPC. The conductance of the lipid bilayer was determined by measuring transmembrane current at a constant transmembrane voltage (*V* = 50 mV). The subsequent calculations were performed assuming that the membrane conductance is related to the membrane boundary potential (φ_*b*_), the potential drop between the aqueous solution and the membrane hydrophobic core, by the Boltzmann distribution (40):

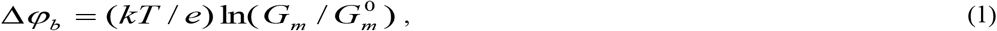

where *G_m_* and 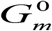 are the steady-state membrane conductance induced by nonactin in the presence and absence of protein, respectively, and *e*, *k* and *T* have their usual meanings. The changes in φ_*b*_ for defined experimental conditions were averaged based on at least three independent experiments (mean ± SD).

### Confocal microscopy of unilamellar vesicles

Giant unilamellar vesicles were formed by the electroformation method on a pair of indium tin oxide slides with using a commercial Nanion vesicle prep pro (Munich, Germany) as previously described (41) (standard protocol, 3 V, 10 Hz, 1 h, 55°C). The lateral phase segregation of membrane components was visualized by an introducing a fluorescent Rh-DPPE probe into the source lipid solution of mixture in chloroform (2 mM): (*1*) 50 mol % DOPC and 50 mol % DPPC; (*2*) 50 mol % DOPC and 50 mol % DMPC; (*3*) 50 mol % DOPC and 50 mol % SM; (*4*) 50 mol % DOPC and 50 mol % TMCL; (*5*) 67 mol % DOPC and 33 mol % CHOL; and (*6*) 40 mol % DOPC, 40 mol% SM and 20 mol % CHOL. Rh-DPPE concentration in the sample was 1 mol %. Rh-DPPE clearly favors liquid disordered phase (*l_d_*) and it is excluded from liquid ordered (*l_o_*) and gel (*s_o_*) phases (42). The obtained liposome suspension was divided into aliquots. An aliquot without protein was used as a control. The experimental samples contained the ratio of lipid to protein of 500:1, 250:1, and 100:1. Vesicles were observed through immersion lenses 60×/1.42 on an Olympus (Germany). The preparations were studied at 25°C. Rh-DPPE was excited by 561 nm light (He-Ne laser). The total number of counted vesicles in sample typically was 10-15. All experiments were repeated three times and the most typical results are presented.

### Differential scanning calorimetry measurements

Differential scanning calorimetry experiments were performed using μDSC 7 EVO microcalorimeter (Setaram Inc., France). Giant unilamellar vesicles were prepared from pure DMPC or pure DPPC by the electroformation method as described above. The obtained liposome suspension contained 4 mM lipid and was buffered by 5 mM HEPES-NaOH at pH 7.4. Protein ApoaCTDH from mM stock solutions in buffer were added to aliquots up to the lipid to protein molar ratio of 500:1, 250:1, 100:1, and 70:1. Liposomal suspension had been incubated with protein for 30 min at room temperature and was heated at a constant rate of 0.2 K·min^-1^. The reversibility of the thermal transitions was assessed by re-heating the sample immediately after the cooling step from the previous scan. At least two independent experiments were done for each system/protein. The temperature dependence of the excess heat capacity was analyzed using Calisto Processing (Setaram Inc., France). The thermal behavior of the liposomes suspension in the absence and presence of the protein was described by the changes in the temperature of the maximum of main phase transition (*T_m_*) and width at half-height of main peak in the endotherm (*T_1/2_*), corresponding to cooperativity of lipid transition.

### Production of liposomes with carotenoids

Canthaxanthin (CAN) and Echinenone (ECN) were extracted from aqueous solutions of COCP and WT OCP by chloroform. For this, a three-fold volume of chloroform was added to a protein solution, vigorously stirred and incubated overnight at + 37 °C. After the incubation, the resulting mixture was centrifuged at 12,000 rpm for 15 min, carotenoid solution in chloroform was carefully removed from the lower part of the test tube. The precipitate was washed with chloroform once again. The resulting carotenoid solution in chloroform (5 ml) was evaporated on a rotary evaporator to a volume of 1 ml. Liposomes with carotenoids were obtained by the method given in the article by Shafaa et al. (43) with slight changes. 100 μl of chloroform solution (20 mg/ml) of egg phosphatidylcholine (Avanti) was added to carotenoid solution in chloroform (1 ml), stirred and evaporated chloroform on a rotary evaporator. The resulting lipid film with carotenoids was solubilized in 500 μl sodium-phosphate buffer (10 μM sodium phosphate, 150 mM sodium chloride, pH 8.0) with subsequent sonication (Finnsonik W-181-T, Ultrasonic, Finland) at a frequency of 40 kHz and a power of 90 watts for 30 minutes. Then the resulting suspension of liposomes was centrifuged at 6000 rpm for 5 minutes for purification from aggregates and carotenoids not incorporated into the liposomes. The supernatant, containing purified liposomes was 4 times filtered through a filter with an average pore diameter of 0.2 μm (Millipore, USA) for standardizing the size of liposomes. The filter was washed with another 100 μl of sodium phosphate buffer and combined with filtered suspension of liposomes. The resulting liposomes were stored in the dark at + 4 °C under argon atmosphere.

### Absorption measurements

Absorption spectra were recorded using a MayaPro2000 spectrophotometer (Ocean Optics, USA). In order to compensate for the effect of light scattering, an integrating sphere BIM-3003 (BroLight, China) was installed in front of the sample. The kinetics of carotenoid transfer was measured as the change of optical density at 550 nm with 100 ms time resolution, the precision of optical density measurement was 5·10^-3^. The temperature of the samples was stabilized by a Peltier-controlled cuvette holder Qpod 2e (Quantum Northwest, USA) with a magnetic stirrer. To estimate the efficiency of the carotenoid transfer under the given conditions, spectral decomposition using reference spectra of carotenoid donors and carotenoid acceptors in the 100% holoform was performed using OrigniPro 9.0 software (OriginLab, USA) by fitting the corresponding contributions of the donor and acceptor spectrum to the spectrum obtained at the end of the mixing experiment. All experiments were performed three times and the most typical results are presented.

### Raman spectroscopy measurements

Resonance Raman spectra of carotenoids in proteins or in liposomes were obtained under continuous excitation at 532 nm. Laser beam was focused on a 0.1 mm glass capillary with a sample. Raman scattered light was collected and subsequently imaged using a confocal microscope-based system (NTMDT, Ntegra Spectra, Russia). The same system was used for the Raman spectroscopy and imaging of HeLa cells, enriched by carotenoids. Processing of Raman images was performed using Nova (NTMDT, Russia) and ImageJ software. At least 10 different HeLa cells were analyzed. *Figure 4B* represents characteristic overlay of Raman signature intensity with the image of HeLa cell in transmitted light.

### Delivery of carotenoids into mammalian cells by CTDH

Cell lines HeLa (derived from cervical cancer cells) were cultured in complete DMEM medium (Thermo Scientific, USA) with 10% (v/v) heat-inactivated fetal calf serum (Thermo Scientific, USA), 100 U/ml penicillin/streptomycin (PanEco, Russia) in a humidified atmosphere with 5% CO_2_ at 37°C. For confocal Raman microscopy, HeLa cells were cultured on glass bottom dishes (POC-R2 Cell Cultivation System, PeCon GmbH, Germany) overnight in 5% CO2 at 37°C. The cells were incubated in a growth medium, containing 1 μM CTDH ECN for 2 hours at 37°C.

The distribution of the TagRFP-CTDH chimera after incubation of cells was recorded using an LSM-710 laser scanning confocal microscope (Carl Zeiss Microscopy, Jena, Germany). Fluorescence was exited at 561 nm, the emission was detected between 565 and 730 nm. The 63x oil Plan-Apochromat objective with numerical aperture of 1.4 was used to obtain high-quality images.

Fluorescence lifetime images were recorded using a FLIM module connected to the LSM-710-NLO (Becker&Hickl, Germany), consisting of a time-correlated photon counting system (TCSPC) SPC-150, hybrid photodetector GaAsP HPM-100-07, SPC-IMAGE software. The TagRFP-CTDH chimera was two-photon exited at 1050 nm by femtosecond laser (Cameleon Ultra II, Coherent, USA).

### Antioxidant activity of carotenoids delivered into mammalian cells by CTDH

Assessment of ROS was performed using dihydroethidium (DHE) (Sigma-Aldrich, D7008) and 2′,7′-Dichlorofluorescin diacetate (DCFDA) (Sigma-Aldrich, D6883), indicators of hydroxyl, peroxyl, superoxide, and other reactive oxygen species (ROS) activity within the cell, according to the manufacturer’s protocol. Experiments were carried out using FACS Canto II (Becton Dickinson) flow cytometer.

To analyze the antioxidant effect of carotenoids we use neuroblastoma cell line Tet21N and antimycin A (AMA), as an inhibitor of electron transport in mitochondria, that has been used as a reactive oxygen species (ROS) generator in biological systems. AMA inhibits succinate oxidase and NADH oxidase, and also inhibits mitochondrial electron transport between cytochrome *b* and *c* (31). This inhibition causes collapse of the proton gradient across the mitochondrial inner membrane and production of ROS.

## Acknowledgments

The authors acknowledge the support of the Russian Science Foundation (Grant no. 18-44-04002), the German Ministry for Education and Research (BMBF grant no. 01DJ15007) and the German Research Foundation (DFG grant no. FR1276/5-1). The study was partially supported by the Russian Foundation for Basic Research grant (no. 18-04-00691 to N.N.S. and no. 19-29-04087 to A.V.S.). Protein–protein interactions were studied in the framework of the Program of the Ministry of Science and Higher Education of Russia.

## Author contributions

E.G.M. conceived the idea and supervised the study; N.N.S., A.V.Z., S.S.E. and E.G.M. designed the experiments; N.N.S., Y.B.S., A.V.Z., E.Y.P., T.A.S., P.A.B., S.S.E., O.S.O., A.V.S., A.V.R. and E.G.M. performed the experiments; N.N.S., S.S.E., T.F., and E.G.M. analyzed the data; E.G.M. and N.N.S. wrote the paper with an input from other authors.

## Data and materials availability

All data is available in the main text or the supplementary materials.

## Supplementary Information

### Spectroscopic characterization of holo-CTDH species

In contrast to OCP-CTD dimer (also called COCP), almost exclusively binding CAN (29, 44), Apo-CTDH may bind both ECN and CAN, which confers different spectral properties and colours to the corresponding holoprotein forms (24). To monitor the dynamics of the holoform assembly or kinetics of the carotenoid transfer, we first studied spectroscopic signatures of ECN and CAN in different environments in more detail (*Figure S1*). CAN bound to a CTDH dimer shows the largest red shift of the absorption maximum among all OCP species. The S_0_-S_2_ absorption maximum located at ~ 560 nm is red-shifted by ~ 10 nm compared to the corresponding transition of CAN in the non-natural OCP-CTD dimer (COCP) from *Synechocystis* (29, 44). The characteristic ~30-nm difference in the position of S_0_-S_2_ absorption maximum of CAN- and ECN-containing CTDH samples has been reported recently (24). This indicates that CAN has a higher conjugation length in CTDH, likely due to the presence of keto groups in each of two β-rings, which both could potentially be conjugated with the polyene chain and involved in hydrogen bonding with Tyr-27 and Trp-110 residues (Tyr-201 and Trp-288 in *Synechocystis* notation), while ECN has only one keto-oxygen. We assume that the nature of the great bathochromic shift of CAN absorption in CTDH is due to hydrogen bonding since mutation of the critical Trp in COCP caused a similar blue shift of CAN absorption (*Figure S1A* and (44)). Absorption of ECN and CAN in liposome membranes is blue-shifted by ~90 nm compared to CTDH holoproteins, and almost identical to the absorption of these carotenoids in organic solvents (45).

**Fig. S1.**
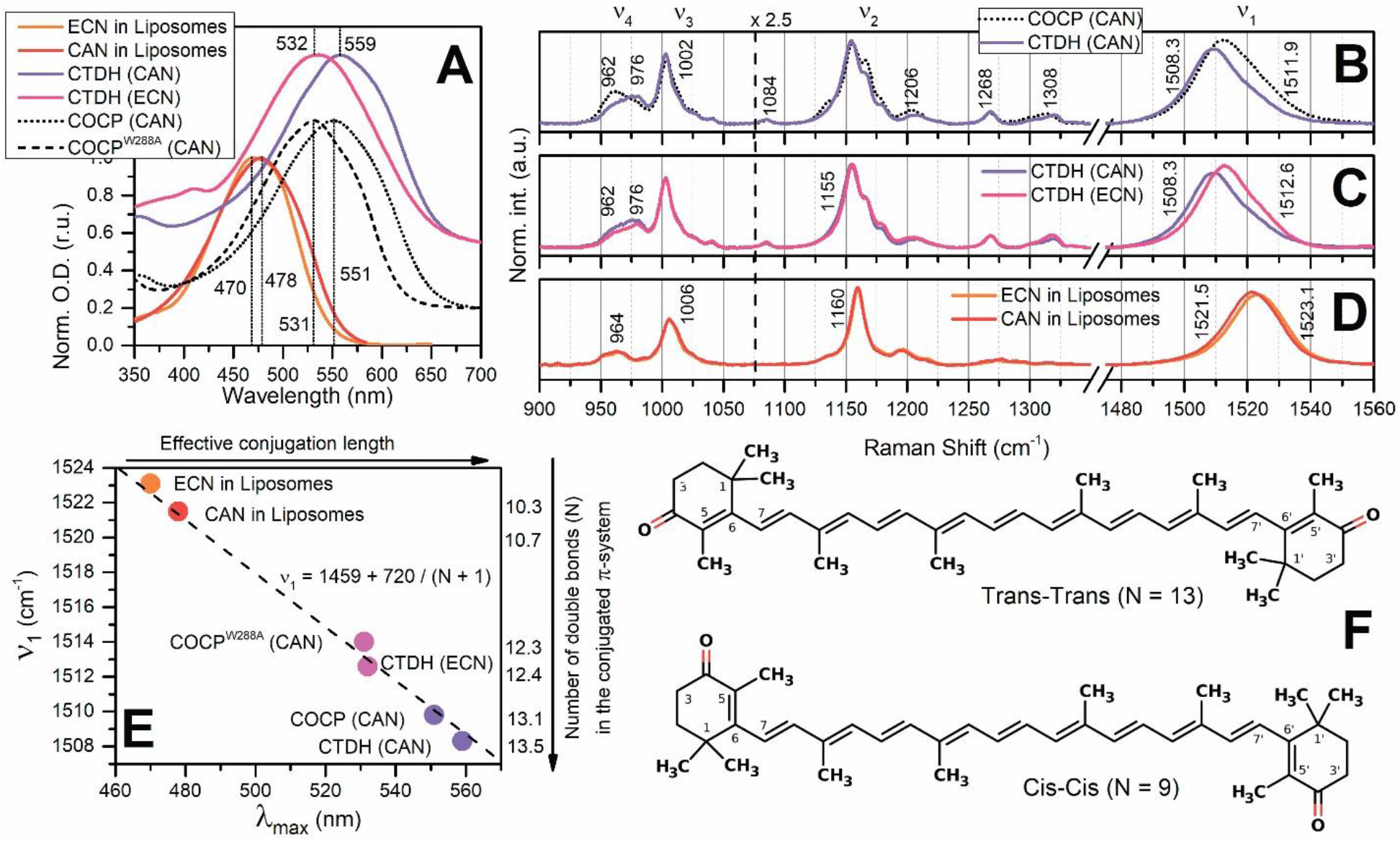
Spectral properties of ECN and CAN in different environments. (**A**) – normalised absorption of CTDH containing CAN (violet) and ECN (purple). Orange and red lines show the absorption of ECN and CAN in liposomes, respectively. Dotted and dashed lines show the absorption of COCP and the W288A mutant thereof, respectively, both containing CAN. Spectra were shifted along the y-axis for the better presentation. (**B** – **C**) Resonance Raman spectra of CTD-like carotenoid proteins recorded with 532 nm excitation. (**B**) – CTDH (solid violet line) and COCP (dotted line), both containing CAN. (**C**) – CTDH containing CAN (violet) and ECN (purple). (**D**) – Raman spectra of ECN in liposomes. The Raman signal in the 900–1060 cm^-1^ region was magnified 2.5 times for the better presentation. Note the x-axis break from 1350 to 1475 cm^-1^. (**E**) – dependency of Raman shift of -C=C-stretching mode (ν_1_) on the position of S_0_-S_2_ maximum (*λ_max_*) in absorption spectra of ECN and CAN in liposomes and proteins. Arrows indicate how the effective conjugation length and the average number of double bonds changes among samples. The average number of conjugated double bonds was calculated using the empirical dependency (46). (**F**) – chemical structures of Cis-Cis and Trans-Trans CAN isomers.

In order to further assess the differences in configurations of ECN and CAN in CTDH and membranes, we used resonance Raman spectroscopy. Although the sequence and secondary structure of COCP and CTDH are very similar, we must consider the fact that the environment for CAN is not exactly the same, since Raman spectra are different. In addition to the already discussed differences of S_0_-S_2_ absorption, differences in conjugation length were estimated by the position of the ν_1_ Raman band. Given the experimentally observed ν_1_ positions and the empirical formula 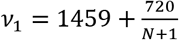,(46), the number of conjugated double bonds (*N*) of CAN changes from 10.7 in membranes to 13.5 in CTDH, which means that in CTDH, all 13 double bonds, including -C=C- and -C=O in the β-rings, are likely coplanar with the polyene chain. In contrast, the β-rings must be out of conjugation when the carotenoid is embedded in a lipid membrane. Thus, we postulate that the transition of the carotenoid from a membrane into the protein is accompanied by a rotation of the β-rings, which results in an increase of conjugation length and thus leads to a characteristic optical response. Since our goal is to study transfer reactions of carotenoids between lipid membranes and proteins and because there is a linear dependence of the absorption *λ_max_* and the shift of the ν_1_ Raman band, we can use any of these spectral characteristics to infer the state of the carotenoid.

The similarity of the spectroscopic properties of CTDH (ECN) and COCP^W288A^ (CAN) indicates that the absence of hydrogen bonds in one of the CTDH subunits (coordinating one of the ECN β-rings, which does not carry a keto group) is equivalent to the absence of two H-bonds with Trps in each of the COCP subunits, resulting in a decrease of the number of conjugated double bonds by ~ 1, comparing to CTDH or COCP with CAN. Such an estimation of the conjugation length changes indicates that even in the absence of hydrogen bonds in one of the CTDH (ECN) subunits protein affects conformation of β-ring, which is at least partially in plane with the polyene chain.

### Echinenone uptake from liposomes

**Fig. S2.**
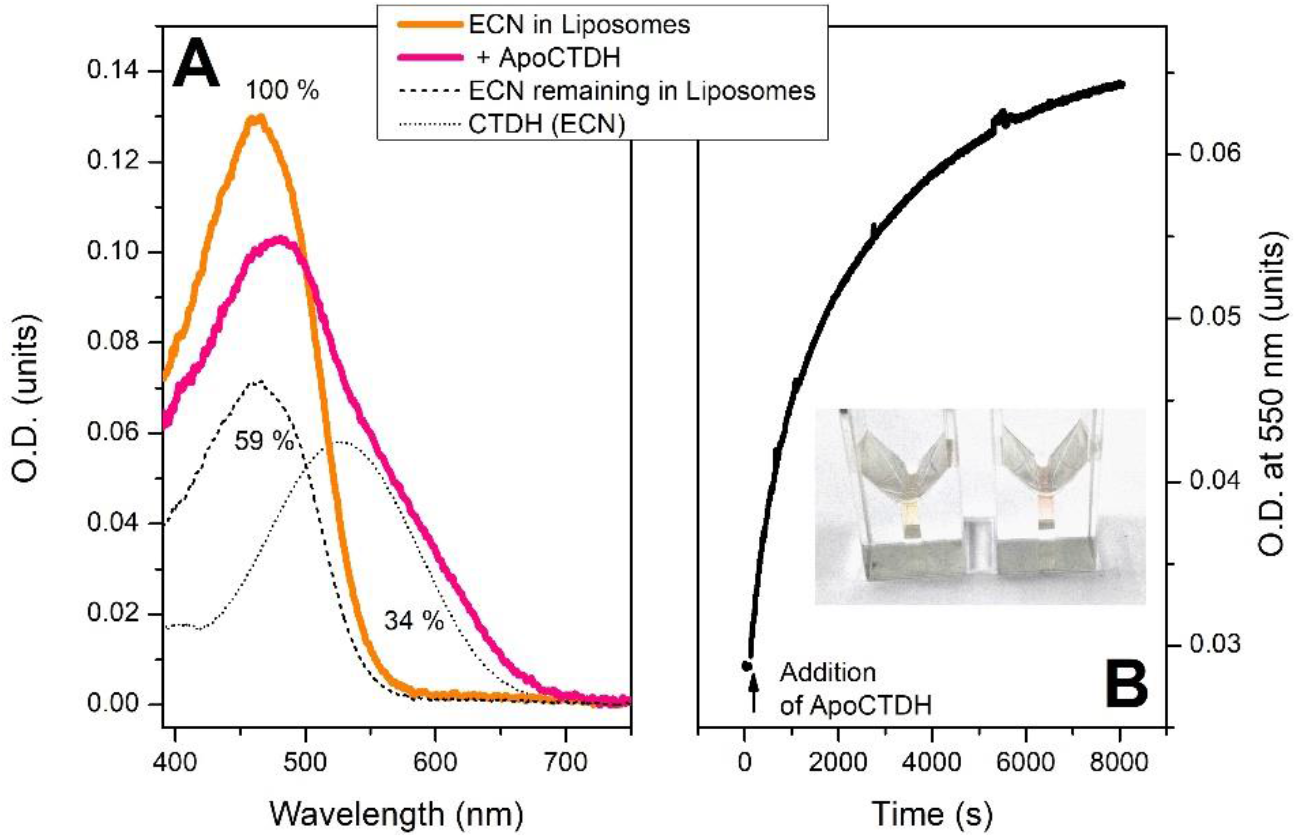
Echinenone uptake from liposomes by ApoCTDH. (A) - absorption spectra of ECN in liposomes before (orange) and after (pink) four hours of incubation with 300 μM of ApoCTDH at 30 °C. Resulting spectrum (pink) was decomposed into components which correspond to absorption of ECN remaining in liposomes (dashed line) and in CTDH (dotted line). Percentiles indicate ECN content in different fractions comparing to indicial concentration in liposomes. Part of carotenoid (~ 7%) was lost due to photodegradation upon the continuous absorption measurements. (**B**) – time-course of accumulation of CTDH (ECN) due to carotenoid uptake from liposomes monitored as increase of optical density at 550 nm. Inset shows cuvettes with solution of ECN-containing liposomes before (left) and after incubation with CTDH.

### ECN delivery from CTDH to liposomes studied by size-exclusion spectrochromatography

**Fig. S3.**
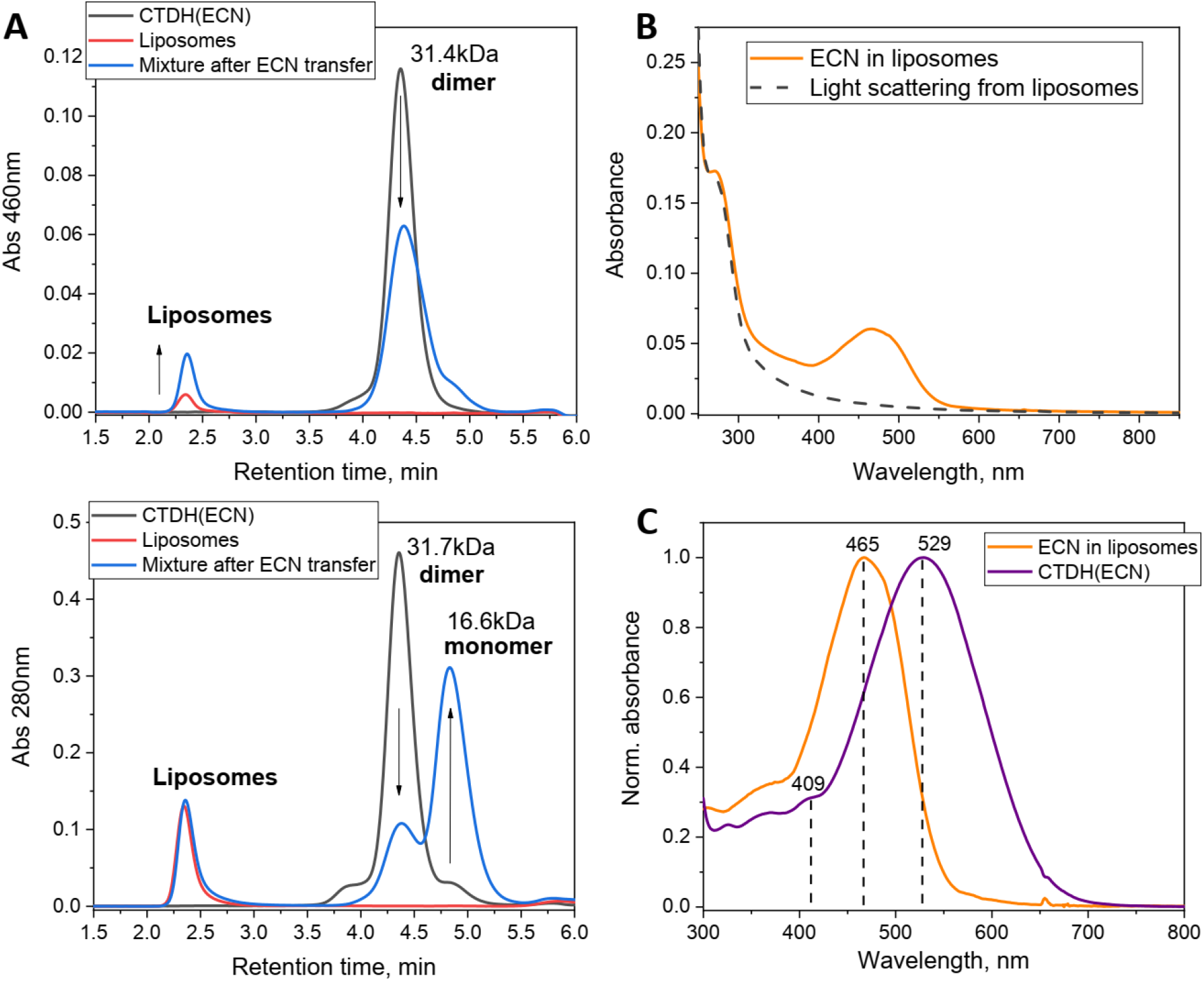
Carotenoid delivery from CTDH to liposomes studied by size-exclusion spectrochromatography. (**A, B**) Elution profiles of CTDH(ECN), liposomes, or their mixture pre-incubated for 40 min at 30 °C to ensure carotenoid transfer followed at either 460 nm or 280 nm on a Superdex 200 Increase 5/150 column (GE Healthcare) operated using a Varian ProStar 335 Diode-Array system with the full spectrum absorbance detection at a 0.45 ml/min flow rate. Apparent Mw for the protein peaks was estimated from column calibration using protein standards (indicated in kDa). Note that liposomes elute in the void volume and give significant light scattering. Arrows indicate the observed changes as the result of carotenoid transfer from CTDH to liposomes accompanied by CTDH(ECN) dimer dissociation. Note that protein does not migrate to the liposome fraction, indicating the absence of tight binding. The most typical result is shown. (**C**). Retrieving the ECN absorption spectrum taking into account light scattering from sample containing pure liposomes in the absence of ECN. (**D**). Absorbance spectra of the initial CTDH(ECN) preparation and ECN in liposomes in the end of the transfer normalized to 0,1.

